# Layilin Regulates Treg Motility and Suppressive Capacity in Skin

**DOI:** 10.1101/2024.12.30.630730

**Authors:** Victoire Gouirand, Sean Clancy, Courtney Macon, Jose Valle, Mariela Pauli, Hong-An Troung, Jarish Cohen, Maxime Kinet, Margaret M. Lowe, Samuel J. Lord, Kristen Skruber, Hobart Harris, Esther Kim, Isaac Neuhaus, Karin Reif, Ali A. Zarrin, Dyche R. Mullins, Michael D. Rosenblum

## Abstract

Regulatory T cells (Tregs) are essential for maintaining immune tolerance in both lymphoid and non-lymphoid tissues. We discovered that layilin, a C-type lectin receptor, is predominantly expressed on Tregs in skin. Layilin was highly expressed on a subset of clonally expanded ‘effector’ Tregs in both healthy and psoriatic skin. Layilin expressing Tregs exhibited a transcriptional profile indicative of enhanced adhesion. Deletion of layilin in Tregs *in vivo* resulted in significantly attenuated skin inflammation. Mechanistically, layilin enhanced Treg adhesion *via* modulation of LFA-1, resulting in distinct cytoskeletal alterations consistent with enhanced focal adhesion and lamellipodia formation. Taken together, we define layilin as a critical regulator of Treg suppressive capacity through modulating motility and adhesion in a non-lymphoid tissue.

## Introduction

Regulatory T cells (Tregs) play a crucial role in mediating peripheral immune tolerance. Although present in lymphoid organs, specialized subsets of these cells stably reside in non-lymphoid tissues, including skin, gut and lung (*1*, *2*). Subsets of Tregs in skin play a crucial role in suppressing hair follicle autoreactivity (*3*). In addition, these cells facilitate hair follicle cycling (*4*), augment wound healing (*5*) and establish and maintain tolerance to skin commensals (*6*, *7*). Many autoimmune and chronic inflammatory diseases are thought to result from an imbalance in the relative abundance and functional state of proinflammatory and regulatory cells. Thus, elucidating the fundamental mechanisms these cells utilize in both healthy and inflamed tissues is crucial in attempts to develop novel therapies aimed at restoring tissue immune homeostasis.

Layilin is a cell surface C-type lectin receptor (*8*, *9*). This family of receptors plays a role in many cellular functions, including adhesion and cell signaling (*10*). Previously, we and others have shown that layilin is preferentially expressed on exhausted CD8+ T cells infiltrating human and murine tumors (*11–13*). In addition, intra-tumoral Tregs express high levels of layilin (*14–16*). Mechanistically, we have shown that layilin functions to anchor Tregs in non-lymphoid tissues, and in doing so, limits their suppressive capacity (*15*). On CD8^+^ T cells, layilin co-localizes with integrin αLβ2 (LFA-1), enhancing its adhesive properties (*13*). Despite emerging evidence that layilin plays an important role in the biology of Tregs in non-lymphoid tissues, little is known about the potential role it plays in autoimmune disease. In addition, whether layilin functions to modulate LFA-1 activation on Tregs remains to be determined. Here, we interrogate the functional biology of Treg expression of layilin in inflammatory skin disease.

## Results

### Layilin is expressed on activated ‘effector’ Tregs in human inflammatory skin disease

To begin to understand the expression pattern of layilin on immune cells in chronic inflammatory skin diseases, we performed single-cell RNA sequencing of CD45^+^ cells sorted from healthy or psoriatic skin. Consistent with previous observations (14), the main population expressing *layilin* are Tregs (**Fig. 1A-C**). There were no changes in the relative expression of *layilin* per cell when comparing Treg clusters from psoriasis skin to healthy control skin (**Fig. S1A**), suggesting that *layilin* expression is not increased on Tregs in inflamed skin. Because layilin is expressed on ∼40-50% of Tregs in both healthy and inflamed skin (*15*), we sought to understand how layilin expressing Tregs differed from layilin non-expressing Tregs at the transcriptional level. To this end, we subclustered Tregs from psoriatic skin and performed pseudo-bulk analysis based on *layilin* gene counts (threshold greater than 0 considered positive for *layilin* expression) (**Fig. 1D&1E**). We found that *layilin* expressing (LAYN>0) and non-expressing (LAYN<0) cells are relatively similar with equivalent levels of *FOXP3* and *CTLA-4* expression (**Fig. 1F**). Similarly to the observation made in **Figure S1A**, we noticed most differences are driven by *layilin* expression (**Fig. S1B**). Focusing on psoriatic skin, we found that 636 genes were significantly differentially expressed (DEGs) between *layilin*-positive and *layilin*-negative Tregs (**Fig. 1G**). Interestingly, several *TNFRSF* family genes and several adhesion molecules, including integrin coding genes (*ITGB7* and *ITGB6*) or cell-cell adhesion (*CADM1*) were expressed at significantly higher levels in *layilin*-expressing Tregs (**Fig. 1G**). Gene set enrichment analysis (GSEA) on DEGs, revealed pathways for T cell activation and cell migration and adhesion to be enriched in *layilin*-expressing Tregs (**Fig. 1H**).

**Figure 1.**
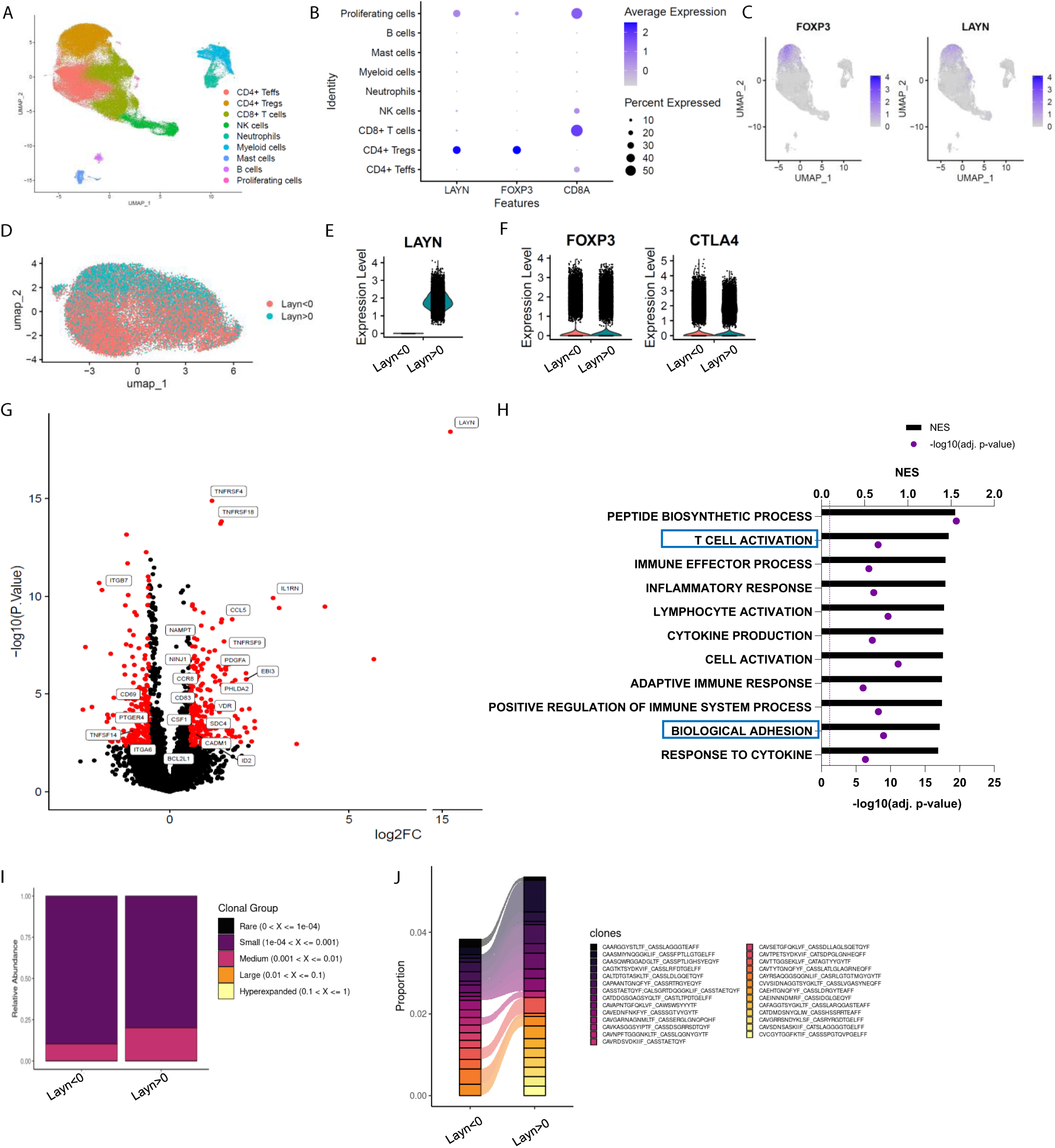
Layilin is preferentially expressed on Tregs in skin and correlates with activation and motility signatures. **A-C.** Single cells RNA-seq of FACS-purified CD45^+^ cells from healthy and psoriatic skin. n= 5 healthy skins and five psoriatic skins. **A.** Representative UMAP showing all clusters found. **B.** Dot plot of *LAYN, FOXP3,* and *CD8A* expression by clusters. **C.** UMAP of *FOXP3* and *LAYN* expression in skin. **D.** UMAP of Tregs subset showing in red *layilin* non-expressing Tregs (LAYN<0) and blue *layilin* expressing Tregs (LAYN>0) based on gene count > 0. **E-F.** Violin plot of *LAYN* (**E**)*, FOXP3* and *CTLA4* (**F**) in subset Tregs from skin followed by resclustering based on *LAYN* gene count > 0. **G.** Volcano plot of *layilin* expressing Tregs (right) compared to *layilin* non-expressing Tregs (left) obtained from pseudo bulk analysis performed on Tregs subset. **H.** Gene set enrichment analysis(GSEA) of top enriched pathways in *layilin* expressing Tregs compared to *layilin* non-expressing Tregs associated with panel F. **I-J.** TCR analysis run on Treg cluster subset by *LAYN* expression. In **I**, we show the relative abundance of TCR clonotype frequencies separated into five groups: rare being expressed between 0 and 10^4^, small between 10^4^ and 0.001, a medium between 0.001 and 0.01, large between 0.01 and 0.1, and hyperexpanded between 0.1 and 1. **J.** Representation of the top expressed clonotypes in subset of Tregs from skin resclustered according to *LAYN* gene count > 0.

T cell receptor (TCR) engagement is a potent inducer of layilin expression on Tregs (*15*). Thus, we hypothesized that layilin expressing Tregs may be specific for tissue antigens and consequently more clonal than Tregs in skin with undetectable layilin expression. To test this hypothesis, we performed TCR repertoire analysis using the same sub-clustering of Tregs as in the pseudo bulk analysis (**Fig. 1I&1J**). Interestingly, *layilin*-expressing Tregs had more expanded clones, with the top 15 expressed clonotypes being more abundant in *layilin*-expressing cells. These findings are similar to previous observations made in CD8^+^ T cells (*13*). Taken together, transcriptional immunophenotyping of Tregs isolated from psoriatic skin suggests that *layilin* expression correlates with more activated, more motile and clonally expanded cells.

### Layilin has minimal effect on Treg activation and suppressive capacity *in vitro*

Activated lymphocytes express receptors and pathways that can either promote or attenuate their effector functions (*17*). In addition, *ex vivo* activation of lymphocytes can induce molecular pathways that only play a meaningful biologic role when the cells are in their natural *in vivo* setting. Thus, we sought to determine if layilin expression on Tregs plays a role in promoting their activation state or suppressive capacity *in vitro*. To do so, we isolated Tregs from peripheral blood of healthy volunteers and expanded them *ex vivo* with anti-CD3 and anti-CD28 coated beads and high dose IL-2. Expanded cells were specifically edited for *layilin* using a well-established CRISPR-Cas9 electroporation protocol (*18*) (**Fig. 2A**). Four days post-electroporation, we consistently observed a maximal reduction in *layilin* expression by ∼30-50% compared to single guide-treated control cells (**Fig. 2B**). After 4 days of suboptimal activation post-electroporation, cells were rested and cultured with or without TCR restimulation. Subsequently, the cells underwent spectral flow cytometry analysis using a detailed panel of Treg activation markers. In these experiments, deletion of layilin did not influence Treg activation across multiple donors with or without TCR stimulation (**Fig. 2C**). To determine if deleting layilin on Tregs impairs their suppressive function *in vitro*, we performed standard Treg suppression assays with cells edited for *layilin* or control guide RNAs. In these experiments, deleting layilin on 30-50% of Tregs had no effect on their ability to suppress either CD4+ or CD8+ T cell proliferation (**Fig. 2D**). Taken together, this data suggests that layilin plays a minimal role, if any, in Treg activation and suppressive capacity *in vitro*.

**Figure 2.**
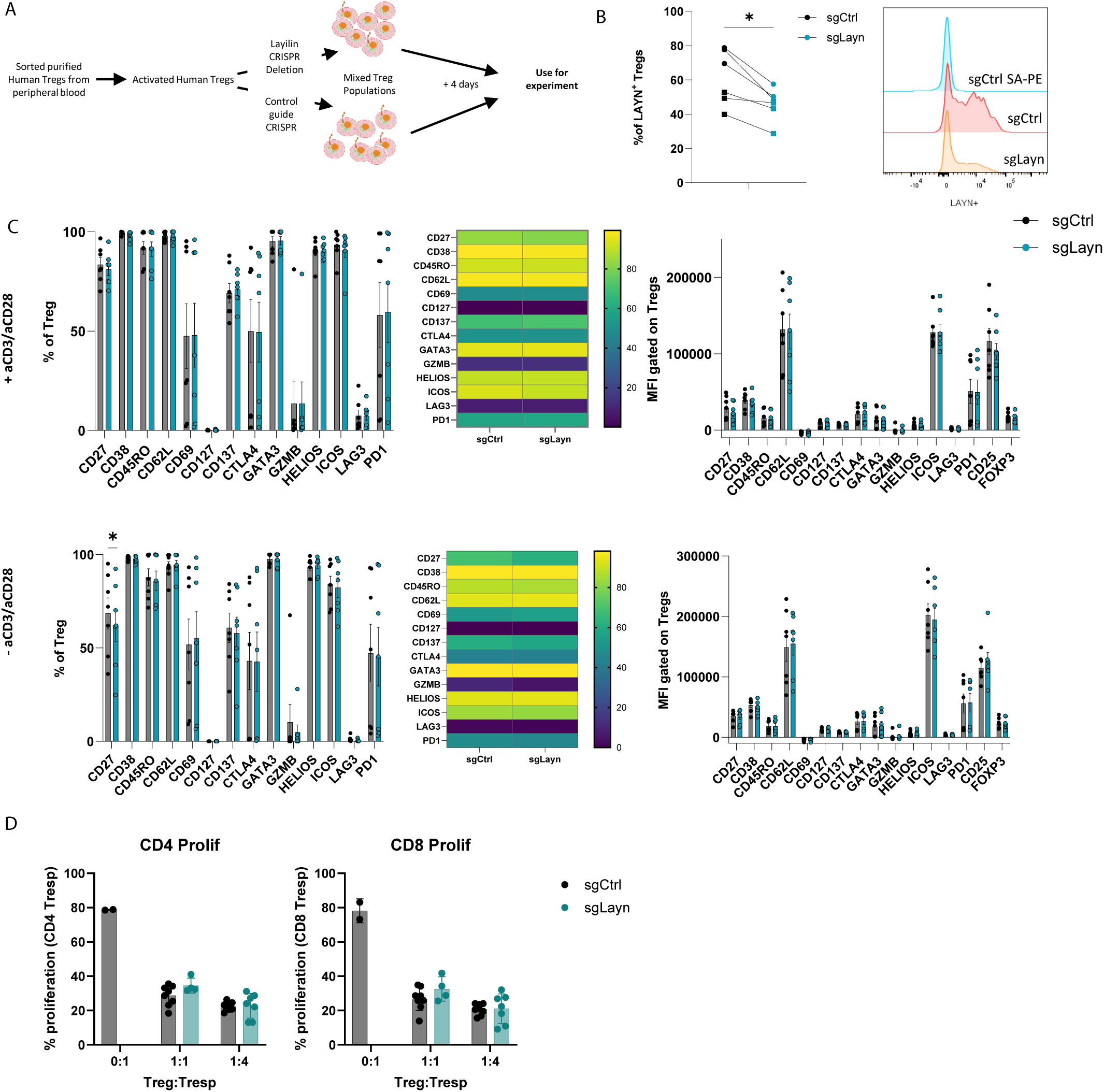
Layilin does not affect Treg activation or suppression *in vitro*. **A.** Schematic experimental design strategy starting with the isolation of human Tregs from peripheral blood, activated and expanded using anti-CD3/anti-CD28 stimulation and IL-2 for 12 days. We deleted *LAYN* using CRISPR/Cas9, and after four days, cells were used for the experiment. **B.** Frequency of LAYN^+^ Tregs gated on Live CD8^-^CD4^+^CD25^+^FOXP3^+^ cells, and representative expression of layilin. n = 6 donors from two independent experiments. Significance was defined by a two-tailed Student’s t-test. *p < 0.05. **C.** Frequencies (on the left) and Mean Fluorescence Intensity (MFI, on the right) of sg-LAYN and sg-CTRL Tregs gated on Live CD8^-^CD4^+^CD25^+^FOXP3^+^ cells expressing the different activation Treg markers listed on the x-axis. The top row graphs show results from cells stimulated with anti-CD3/anti-CD28; the bottom graphs show results obtained from cells without anti-CD3/anti-CD28 stimulation. n = 6 donors from two independent experiments. Significance was defined by using a multiple-paired t-test. **D.** Proliferation percentage measured using cell trace violet (CTV) for CD4^+^ T responder (left panel) or CD8^+^ T responder (right panel) after three days of suppression assay. Results are presented in the x-axis, and the different ratios of sg-LAYN and sg-CTRL Tregs and T responders used for the assay represent two independent experiments. Significance was defined by two-way ANOVA followed by Tukey’s multiple comparisons test.

### Layilin attenuates Treg suppression *in vivo*

We have previously shown that layilin acts to anchor Tregs in tissues and that deletion selectively in these cells results in enhanced tumor growth (*15*). Thus, despite having limited to no role in Treg activation and suppression *in vitro* (**Fig. 2**), we hypothesized that layilin expression on Tregs *in vivo* would attenuate their suppressive capacity. To test this hypothesis, we utilized the well-established imiquimod (IMQ) model of skin inflammation (*19*, *20*), where Tregs are known to play a significant role (*21–23*). To inducibly and selectively delete *layilin* in Tregs, we crossed FOXP3-ERT2-Cre mice (Foxp3^CRE^) (*24*) with Layn^fl/fl^ mice (generated previously by our lab (*13*)). To delete layilin in Tregs, the resultant Foxp^CRE^/Layn^fl/fl^ mice were treated with systemic tamoxifen from 6 days to 2 days prior to beginning IMQ application on dorsal back skin for 6 days (**Fig. 3A**). Because there are no working antibodies that reliably detect murine layilin by flow cytometry, we performed qRT-PCR on sort-purified Tregs from Foxp^CRE^/Layn^fl/fl^ and Layn^fl/fl^ mice after tamoxifen treatment to assay for *layilin* deletion. These experiments confirmed a significant reduction in *layilin* expression in Tregs from Foxp^CRE^/Layn^fl/fl^ mice (**Fig. S2A**). Clinical signs of disease, including skin lesions, scaling, and erythema were quantified in a blinded fashion throughout the duration of the IMQ treatment. Consistent with our hypothesis, we observed a significant reduction in the clinical severity of skin inflammation in Foxp^CRE^/Layn^fl/fl^ mice treated with tamoxifen compared to age- and gender-matched littermate Foxp^CRE^ control mice treated with tamoxifen (**Fig. 3B & 3C**). Flow cytometric quantification of immune cell infiltrate and cellular cytokine production was performed on day 6. Foxp^CRE^/Layn^fl/fl^ mice had a significant reduction in the absolute number of skin infiltrating CD4+ Teff cells and CD8+ T cells when compared to controls (**Fig. 3D**). In addition, there was a significant reduction in the number of TNFa-producing CD4+ Teff cells, IL-17-producing CD8+ T cells, and IL-17-producing Tregs in Foxp^CRE^/Layn^fl/fl^ mice compared to controls (**Fig. 3E & Fig. S2B**). There were no significant differences in the expression of select Treg activation markers between layilin-deleted and control Tregs (**Fig. 3F & Fig. S2C**).

**Figure 3.**
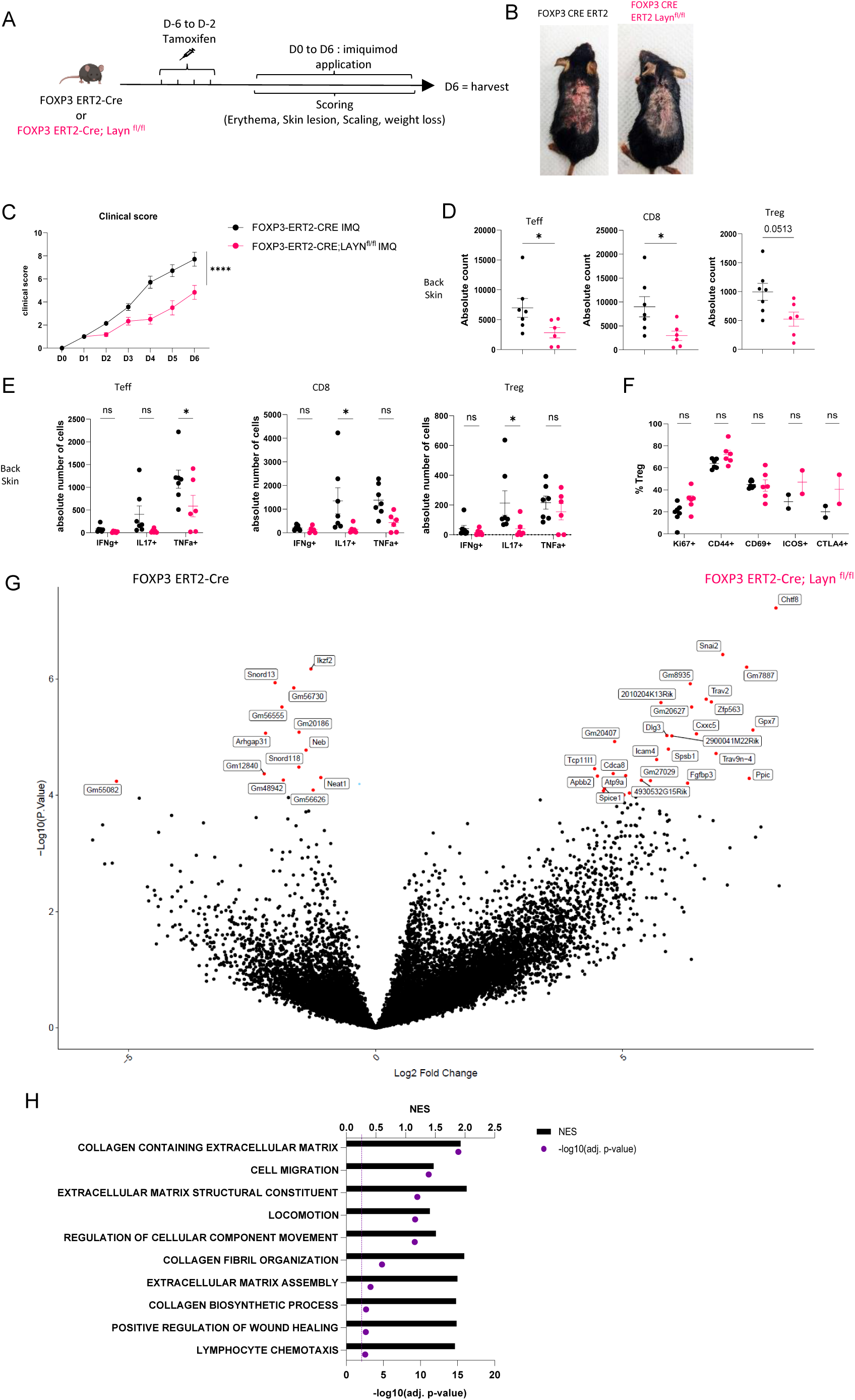
Layilin limits Treg-mediated suppression of inflammation *in vivo*. **A.** Experimental design strategy of the imiquimod (IMQ) model. After four days of tamoxifen injection, mice were rested for two days before the start of the IMQ treatment. They received a daily dose of IMQ on shaved back skin and **were** scored for erythema, scaling, skin lesions, and weight loss. After six days of treatment, the mice were euthanized. **B.** Representative pictures of mouse back skin after five days of treatment from each experimental group. **C.** Overall clinical score over time. n =7 mice from three independent experiments. Significance was defined by two-way ANOVA. ****p<0.0001. **D.** Absolute count of Teff, CD8, and Tregs obtained from each experimental group, respectively gated on Live CD3^+^TCRβ^+^ and CD4^+^FOXP3^-^ or CD8^+^, CD4^+^FOXP3^+^. n =7 and 6 mice per group from three independent experiments. Significance was defined by a two-tailed Student’s t-test. *p < 0.05. **E.** Absolute count of cells Teff, CD8, and Treg either IFNɣ+, IL17+, or TNFα^+^. n =7 and 6 mice per group from three independent experiments. Significance was defined by a multiple-paired t-test. *p < 0.05. **F.** Frequencies of Tregs activation markers gated on Live CD3^+^TCRβ^+^ CD4^+^FOXP3^+^ Tregs. n =7 and 6 mice per group from three independent experiments, except ICOS and CTLA from 2 mice and one experiment. Significance was defined by a multiple-paired t-test. **G.** Volcano plot of differential expression of genes from sorted Tregs gated on GFP^+^ from FOXP3-ERT2-Cre; Layn^fl/fl^ (on the right) or FOXP3-ERT2-Cre (on the left) after four days of treatment. Red dots represent differentially expressed genes; black dots are not differentially expressed ones. n = 5 mice per group from two independent experiments. **H.** Top deregulated pathways obtained from a GSEA comparing FOXP3-ERT2-Cre; Layn^fl/fl^ to FOXP3-ERT2-Cre Tregs. The top axis and bar plot represent the Normalized Enrichment Score (NES). The bottom axis and dot represent the adjusted p-value associated with pathways enrichment. n = 5 mice per group from two independent experiments.

To comprehensively interrogate how layilin deletion alters Treg biology *in vivo* at the transcriptional level, we performed bulk RNA sequencing. RNA was isolated from GFP-expressing Tregs sort purified from back skin after four days of IMQ treatment and subjected to low-input RNA sequencing (**Fig. S2D & S2E**). These experiments identified several DEGs in Foxp^CRE^/Layn^fl/fl^ mice relative to Foxp^CRE^ control mice both treated with tamoxifen (**Fig. 3G**). Interestingly, GSEA analysis of DEGs revealed enrichment in pathways involved in cellular movement, locomotion and migration (**Fig. 3H**). Taken together, these results reveal that layilin attenuates Treg suppression *in vivo* without altering their activation capacity and is consistent with previous studies that suggest that the anchoring function of layilin limits Treg’s ability to optimally mediate immune regulation in skin (*15*).

### Layilin enhanced Treg adhesion in an LFA-1-dependent manner

We have previously shown that layilin co-localizes with LFA-1 on CD8+ T cells and functions to augment LFA-1-mediated adhesion of these cells (*13*). Thus, we set out to determine if layilin functions in a similar fashion on Tregs. In addition, we sought to determine whether layilin expression on Tregs influences motility pathways at the transcriptional level. To this end, we sorted purified Tregs from human peripheral blood, CRISPR-edited these cells for *layilin* (or control guide RNA), sort-purified these cells based on layilin expression, and performed bulk RNA sequencing on highly pure layilin expressing or layilin-edited cells (**Fig. 4A**). Consistent with experiments described above where layilin was deleted in Tregs *in vivo* in mice (**Fig. 3**), several genes involved in cellular locomotion, migration and adhesion were differentially expressed in layilin-expressing vs. layilin-edited cells (**Fig. 4B & 4C**).

**Figure 4.**
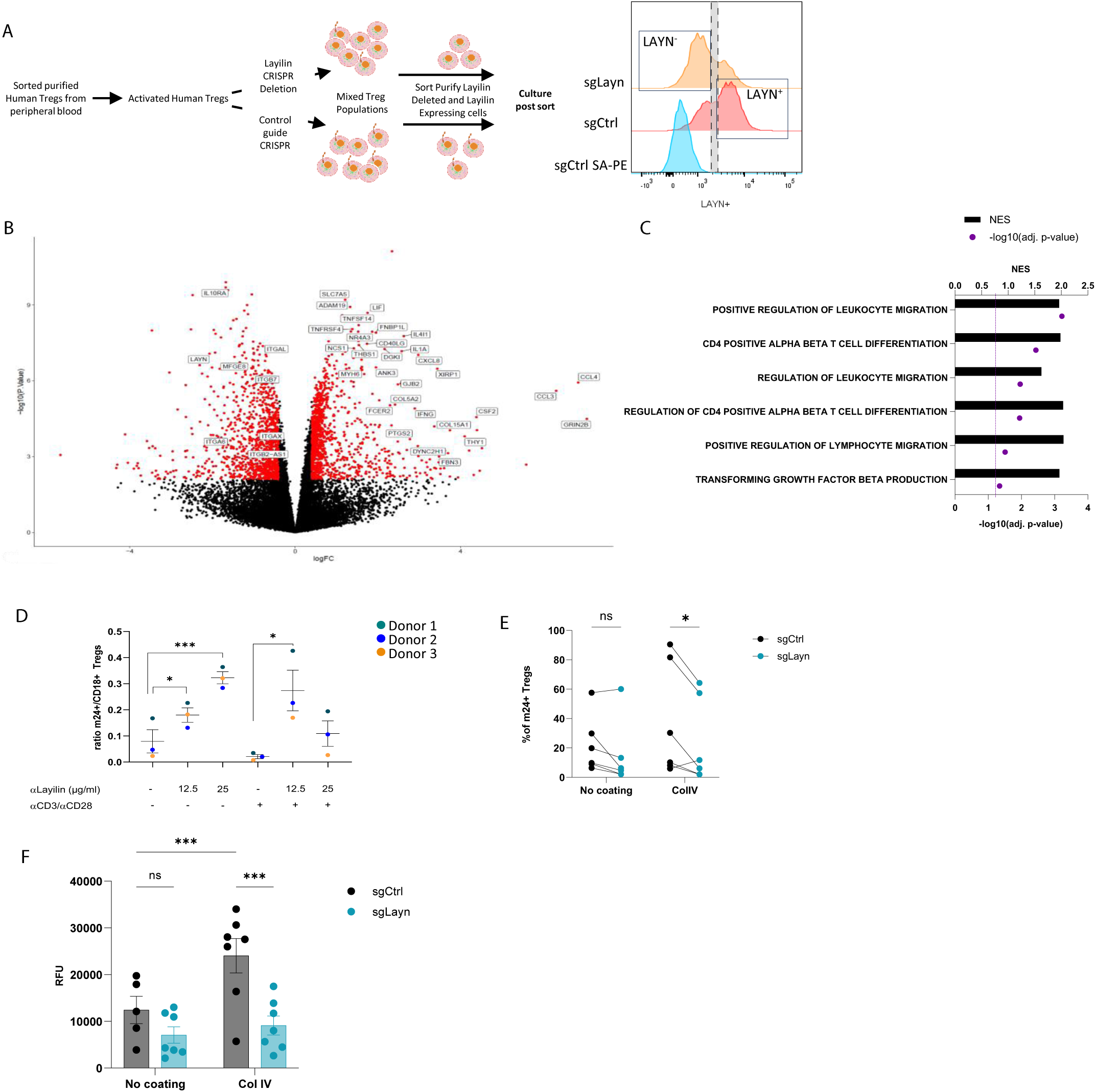
Layilin enhances Treg adhesion and LFA-1 activation. **A.** The experimental design strategy starts with the isolation of human Tregs from peripheral blood, which were activated and expanded using anti-CD3/anti-CD28 stimulation and IL-2 for 12 days. We deleted *LAYN* using a CRISPR/Cas9; after four days, cells were sorted by flow cytometry based on LAYN expression, as represented by the representative histogram of LAYN expression on sgCtrl (stained with or without primary antibody detecting layilin followed by streptavidin-PE staining) or sgLayn Tregs, to obtain a pure population of LAYN^+^ or LAYN^-^ Tregs. After sort, cells were placed in culture for follow-up experiments presented in panels B, C and F. **B.** Volcano plot of differential genes expressed in LAYN^+^ (on the left) or LAYN^-^ (on the right) Tregs put in culture for 24 hours in the presence of anti-CD3/anti-CD28. Red dots represent differentially expressed genes; black dots are not differentially expressed ones. n = 4 paired donors for each group. **C.** Top deregulated pathways obtained from a GSEA analysis comparing LAYN^+^ to LAYN^-^ Tregs. The top axis and bar plot represent the Normalized Enrichment Score (NES). The bottom axis and dot represent the adjusted p-value associated with pathways enrichment. n = 4 paired donors for each group. **D.** Ratio of m24^+^ to total CD18^+^ Tregs gated on Live CD8^-^CD4^+^CD25^+^FOXP3^+^ cells after three days culture with or without anti-CD3/anti-CD28 stimulation and in the presence of a layilin crosslinking antibody as described on the x-axis for 20 minutes during LFA-1 activation staining. n = 3 donors. Significance was defined by one-way ANOVA followed by Dunnet’s multiple comparisons test. *p < 0.05, ***p<0.001**. E.** Frequency of m24^+^ sg-LAYN and sg-CTRL Tregs gated on Live CD8^-^CD4^+^CD25^+^FOXP3^+^ cells after 24 hours of culture in a non-coated plate or collagen IV coated plate. n = 6 donors from two independent experiments. Significance was defined by two-way ANOVA followed by Sidak’s multiple comparisons test. *p < 0.05. **F.** Numbers of cells attached to the bottom of the well are measured by the cell titer blue (expressed by relative fluorescence unit). n = 7 donors from two independent experiments. Significance was defined by two-way ANOVA. ns is non-significant, , ***p<0.001.

Lymphocyte Function-associated Antigen-1 (LFA-1) is critical for the development and function of Tregs (*25*, *26*). LFA-1 is present either in a low-affinity inactive form or a high-affinity active form (*27*, *28*). In its active form, LFA-1 mediates binding to ICAM-1 (Intercellular Adhesion Molecule 1) and cell-cell adhesion (*29*, *30*). To test if layilin modulates LFA-1 function on Tregs, we utilized an anti-human layilin crosslinking antibody (*13*). Layilin-expressing human Tregs were treated with anti-layilin crosslinking antibody and LFA-1 activation was quantified by flow cytometry using the m-24 antibody binding, a monoclonal antibody (mAb) that specially recognizes LFA-1 in its open, active conformation (*31*). These experiments were performed with and without anti-CD3/anti-CD28 stimulation to determine whether TCR stimulation can enhance the effect of layilin on LFA-1 activation. Both in non-stimulated and stimulated conditions, cross-linking layilin on Tregs resulted in increased LFA-1 activation, as measured by increased m-24 mAb staining (**Fig. 4D**).

To bolster our findings with layilin cross-linking antibodies, we set out to repeat these experiments with a natural ligand for layilin. Layilin has been shown to bind to collagen IV (*32*). Thus, to confirm layilin’s impact on LFA-1 activation, we used our CRISPR system to reduce layilin expression (**Fig. S3C**) and plated the cells from five different donors on non-coated or collagen IV-coated plates. Consistent with results using layilin cross-linking mAbs, these experiments revealed that CRISPR-mediated *layilin* editing reduces LFA-1 activation only in the presence of collagen IV (**Fig. 4E**). Finally, we tested the direct adhesive capacity of Tregs using an adhesion assay under the same coated plate conditions. Cells were sort-purified on layilin expression (as described above) to obtain pure populations of layilin-expressing and layilin-deficient Tregs. Consistent with our previous findings, Tregs deficient in layilin displayed a reduced capacity to bind to collagen IV coated plates but not uncoated plates (**Fig. 4F**). Taken together, this data reveals that engagement of layilin on Tregs results in enhanced LFA-1 activation and cellular adhesion.

### Layilin induces cytoskeleton changes in Tregs

Actin polymerization is one of the major cytoskeleton changes involved in cell migration and adhesion (*33*). Thus, we hypothesized that signaling through layilin would induce cytoskeleton changes consistent with increased adhesion. To test this, we measured actin dynamics in cells by performing F-actin staining on Tregs sort-purified based on layilin expression after CRISPR-editing with *layilin*-specific or control guide RNAs (as described in Fig. 4A). These experiments were performed in the presence of layilin ligand (collagen IV) or irrelevant ligand (fibronectin). These experiments revealed increased F-actin staining and increased cellular projections (consistent with lamellipodia) in layilin-deficient cells only in the presence of collagen IV (**Fig. 5A & 5B**). As previously shown, layilin contains a talin binding domain, which we have shown to be essential for layilin-LFA-1 interaction in exhausted CD8^+^ T cells (*13*). Furthermore, talin, vinculin, and paxillin (PAX) are recruited to form a complex and interact with the binding domain of high-affinity conformation integrins, leading to focal adhesion and slowing cells (*34*). Conversely, an increase in polymerized actin content and cellular protrusions correlate with a higher motile profile (*35*, *36*). Lamellipodia are the main protrusive structures formed by T cells during migration and require actin-related protein 3 (ARP3) (*37*). Taken together, we propose a working model (**Fig. 5C**) in which layilin, by its interaction with LFA-1 and talin, stabilizes the formation of focal adhesion complexes and favors a more anchored Treg cell phenotype. Conversely, the absence of layilin promotes cell motility, at least in part, by enhancing lamellipodia formation.

**Figure 5.**
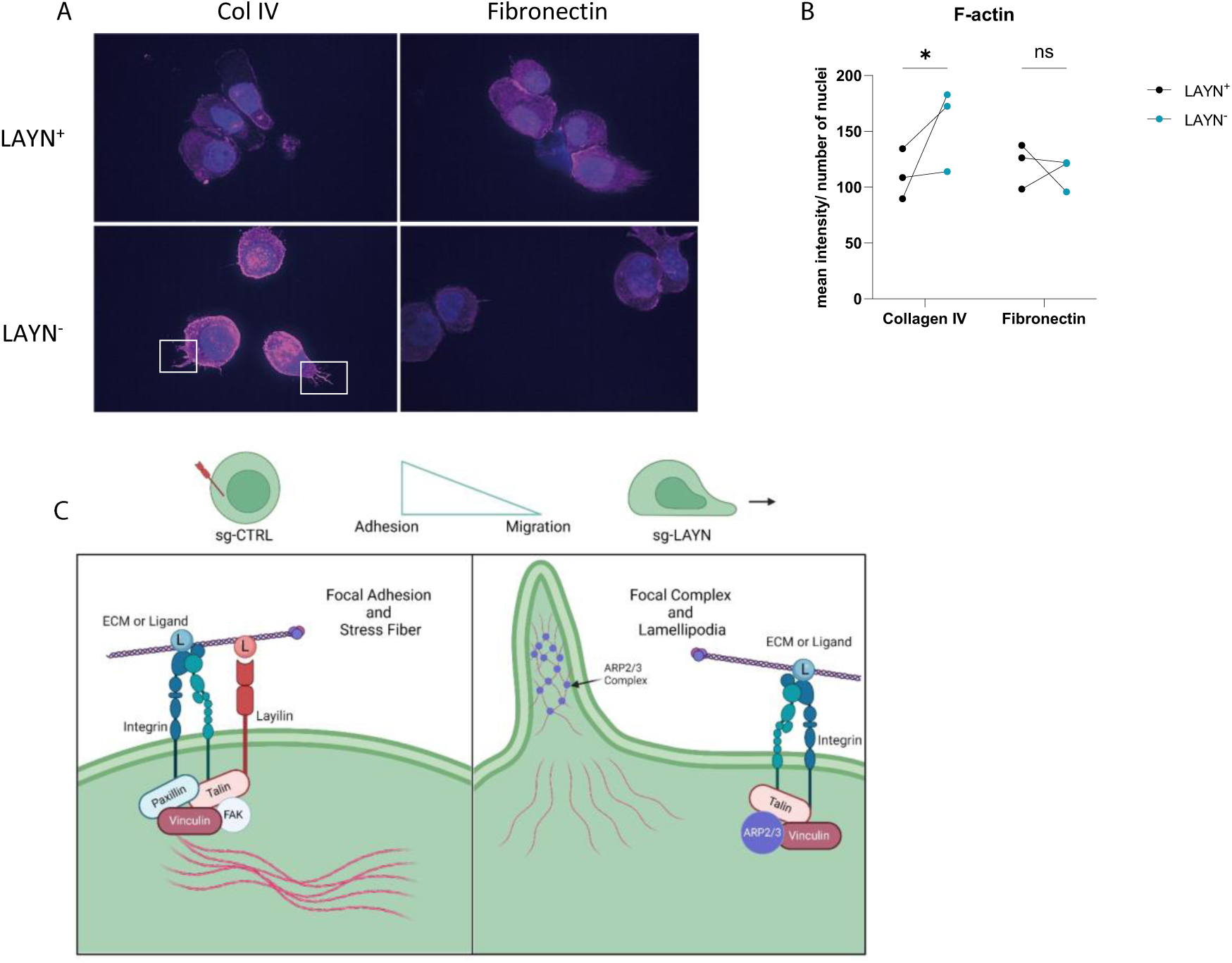
Layilin induces cytoskeleton changes in Tregs indicative of reduced motility. **A.** Representative pictures of LAYN^+^ (on the top) or LAYN^-^ (on the bottom), Tregs phalloidin staining (in pink) and nuclei (in blue) after a 48-hour culture on collagen IV (left) or fibronectin (right) coated plates. White boxes highlight protuberances at the membranes**. B.** Quantification of phalloidin staining associated with pictures in A. n = 3 donors; ten pictures were quantified for each donor, and experiments were repeated independently twice. Significance was defined by two-way ANOVA followed by Sidak’s multiple comparisons test. *p < 0.05. **C.** Graphical hypotheses of layilin pathway mechanism of action. In the presence of layilin, it interacts with LFA-1 and maybe other integrins and stabilizes the formation of a focal adhesion complex, leading to the anchoring mechanism of the cells. Without layilin, integrins are less prone to adhesion mechanisms, and the ARP3 complex may act in lamellipodia formation, leading to more migratory Tregs.

## Discussion

In this study, we discovered that layilin, a C-type lectin cell surface receptor, is highly expressed by Tregs in skin. The summation of our data from results presented herein and previous studies (*15*), suggest that layilin modulates Treg motility in tissues. This effect on motility is mediated, at least in part, through the modulation of LFA-1 activation and contributes to cytoskeletal remodeling necessary for cellular adhesion and movement.

Our data supports a model whereby layilin expression is induced on Tregs upon entering non-lymphoid tissues such as skin. This acts to anchor Tregs in the tissue, possibly adjacent to (or within) collagen IV-rich blood vessels (*38*, *39*). In this capacity, layilin may help to sequester Tregs to a defined niche facilitating their ability to maintain tissue immune homeostasis in the steady state. Conversely, in pathological situations such as tumors and chronic inflammation, the motility of Tregs may play a larger role in their functionality, allowing them to navigate to various tissue microanatomic niches to attenuate specific cell types and inflammatory mediators. Notably, several studies have underscored the crucial role of adhesion properties, particularly those mediated by ICAM-1, the main adhesion molecule expressed by T cells, in enabling effective Treg suppression capacity (*40–42*). In this context, layilin expression on Tregs most likely acts to curtail their suppressive capacity by limiting their motility. Indeed, 2-photon microscopy studies revealed that layilin deficient Tregs are relatively more motile in inflamed skin when compared to layilin expressing control Tregs (*15*). In support of this theory, herein, we show that reducing layilin expression on Tregs results in enhanced immune regulation in a skin inflammation model, as demonstrated by a significant reduction of pro-inflammatory cells, observed in Foxp^CRE^/Layn^fl/fl^ mice compared to controls. In contrast, it is interesting to highlight that, *in vitro*, layilin plays a minimal role in Treg activation and suppressive capacity *in vitro.* This suggests the role of layilin in the suppression capacities of Tregs depends on the more complex, dynamic and collagen rich *in vivo* environment.

An intriguing finding is the association between layilin and LFA-1 activation observed in Tregs, echoing previous discoveries in CD8+ T cells (*13*). LFA-1 is crucial for T cell engagement at the immune synapse (*43*, *44*). While the modulation of LFA-1 by layilin may enhance Treg interactions with other cells, potentially boosting their suppressive abilities, it could also confine their impact to a narrowly defined area due to reduced motility.

Layilin’s role in enhancing the effector functions of CD8+ T cells in tumors (*11*, *13*, *45*) and limiting Treg suppressive capacity in inflamed tissues, suggests that it could be a valuable therapeutic target for autoimmune and chronic inflammatory diseases. A layilin neutralizing anti-body could attenuate CD8+ T cell killing and enhance Treg suppression, effectively working in concert to dampen tissue inflammation. Tissue specific autoimmune diseases thought to be mediated by autoreactive CD8+ T cells and sub-optimally controlled by Tregs would be attractive indications. However, more research is necessary to understand the role of layilin on non-immune cells, such as gastrointestinal epithelial cells (*9*, *46*) to help build confidence around this therapeutic hypothesis.

## Materials and Methods

### Experimental Animals

Foxp3^ERT2-GFPCre^ mice were purchased from The Jackson Laboratory (Bar Harbor, ME) and were bred and maintained in the University of California San Francisco (UCSF) specific pathogen-free facility. As previously described by our group, Layn^fl/fl^ mice were created by inserting LoxP sites flanking exon 4 of the *layilin* gene using CRISPR-Cas9 (*15*). Layilin was deleted specifically on Tregs by crossing Layn^fl/fl^ mice to Foxp3^ERT2-GFPCre^ mice upon treatment with tamoxifen. All mouse experiments were performed on 7–12-week-old animals. All mice were housed under a 12-hour light/dark cycle. All animal experiments were performed following guidelines established by the Laboratory Animal Resource Center at UCSF, and all experimental plans and protocols were approved by IACUC beforehand.

### Mouse Tissue Processing

Briefly, mouse skin was minced and digested in a buffer containing collagenase XI, DNase, and hyaluronidase in complete RPMI in an incubator shaker at 225 rpm for 45 minutes at 37°C. Once the single-cell suspensions were obtained, cells were counted using an automated cell counter (NucleoCounter NC-200, Chemometec). 2-4 x 10^6^ cells were stained, and flow cytometric analysis was performed.

### Human specimens

Normal healthy human skin was obtained from patients at UCSF undergoing elective surgery, and the specimens received were deidentified (study number 19-29608) and certified as Not Human Subjects Research. Blood samples were obtained from healthy adult volunteers (study number 12-09489) or ordered from the Standford Blood Center. Biopsies of psoriasis were obtained with a 6-mm punch biopsy tool (study number 19-29608). For all fresh human tissue samples, active/flaring disease was confirmed both clinically and histologically. Studies using human samples were approved by the UCSF Committee on Human Research and the IRB of UCSF. Informed written consent was obtained from all patients.

### Human Skin Digestion

Skin samples were stored in a sterile container on gauze and PBS at 4°C until digestion. The skin was processed and digested as previously described. Briefly, hair and subcutaneous fat were removed, and skin was cut into small pieces and mixed with digestion buffer containing 0.8 mg/ml Collagenase Type 4 (4188; Worthington), 0.02 mg/ml DNAse (DN25-1G; Sigma-Aldrich), 10% FBS, 1% HEPES, and 1% penicillin/streptavidin in RPMI medium and digested overnight in an incubator. They were washed (2% FBS, 1% penicillin/ streptavidin in RPMI medium), double filtered through a 100-μm filter, and cells were pelleted and counted.

### Tregs are derived from human peripheral blood

CD4^+^ T cells were purified using RosetteSep (Stem Cell) followed by a gradient of Ficoll-Paque Plus (GE Healthcare) from the peripheral blood of healthy volunteers. Once isolated, CD4^+^ T cells were stained and sorted by gating on live CD4^+^CD8^-^CD25^hi^CD127^lo^ cells. Purified Tregs were *ex vivo* expanded for 12 days in complete X-Vivo media (Lonza) with 500U/ml IL-2 (2000U/ml, Tonbo Biosciences) and stimulated with anti-CD3/CD28 beads at cells:beads ratio of 1:1 (Gibco). At D12, cells were frozen for later experiments.

### Human Tregs engineering/CRISPR

To engineer human Treg, we used a CRISPR-cas9 SNP protocol. We thawed D12 Tregs and rested them for 4 hours. Afterward, we mixed 22.2 pmol of Streptococcus pyogenes Cas9 with 88.8 pmol of scramble single guide (sg-CTRL) or *layilin* single guide (sg-LAYN, UUCUCGAA-GACUGAACUUUG) and incubated for 10 minutes at room temperature to form the RNP complex. During the incubation, we harvested the cells, counted them using the NucleoCounter, and resuspended them in P3 buffer cells at a 2 million/20ul concentration. Once the RNP complex was formed, an electroporation enhancer (IDT) was added at a ratio of 1:2 (EE:RNP) to the cells in an electroporation plate. Cells were electroporated using the DG-113 program on a 4D-Nucleofector 96-well unit (Lonza). After four days, efficiency was measured, and cells were used for the experiment.

### Single Cell RNA Sequencing

Sequencing data will be uploaded to the National Center for Biotechnology Information Gene Expression Omnibus (GEO) upon acceptance of the manuscript. scRNA-seq and scTCR-seq libraries were prepared by the UCSF Core Immunology laboratory using the 10X Chromium Single Cell 5′ Gene Expression and V(D)J Profiling Solution kit, according to the manufacturer’s instructions (10X Genomics). 150 paired-end sequencing was performed on a Novaseq 6000 instrument. The Cell Ranger analysis pipelines were then used to process the generated sequencing data. Data were demultiplexed into FASTQ files, aligned to the GRCh38 human reference genome, and counted, and TCR library reads were assembled into single-cell V(D)J sequences and annotations. For gene expression analysis, the R package Seurat was used. Filtered gene-barcode matrices were loaded, and quality-control steps were performed (low-quality or dying cells and cell doublets/multiplets were excluded from subsequent analysis). Data were normalized and scaled, then linear dimensional reduction using principal-component analysis was performed. Highly variable genes were used to perform unsupervised clustering and nonlinear dimensional reduction with UMAP was used to visualize the data.

For differential gene expression analysis comparing LAYN-expressing cells to LAYN-non-expressing cells, data were subsetted into LAYN negative (LAYN<0) and LAYN positive (LAYN>0) using LAYN expression > 0 as a threshold. For TCR analysis, samples were downsampled to get the same number of cells for each donor. All plots were generated using the R packages ggplot2 and cowplot.

### Conventional and spectral flow cytometry

Single-cell suspensions were counted and pelleted. Cells were washed and stained with Ghost Viability dye (Tonbo Biosciences) and antibodies against surface markers in PBS. Cells were fixed and permeabilized for intracellular staining using a FoxP3 staining kit (eBiosciences) and then stained with antibodies against intracellular markers. Fluorophore-conjugated antibodies specific for human or mouse surface, and intracellular antigens were purchased from SinoBiological, Cytek, BD Biosciences, eBiosciences, or BioLegend. The following anti-human antibodies and clones were used: layilin (Clone #07), PE-Streptavidin, CD3 (UCHT1), CD4 (SK3), CD8 (SK1), CD45RO (UCHL1), FoxP3 (PCH101), CD25 (M-A251), CTLA4 (14D3), ICOS (ISA-3), CD27 (O323), CD11c (3.9), HLA-DR (L243), CD137 (4B4-1), PD1 (EH12.2H7), Ki67 (EH12.2H7), LAG3 (11C3C65), CD38 (HIT2), CD62L (DREG56), CD69 (FN50), CD127 (eBioRDR5), GATA3 (L50-823), GZMB (GB11), HELIOS (22F6). Layilin Ab was biotin-tagged using Miltenyi’s One-Step Antibody Biotinylation kit (130-093-385). The following anti-mouse antibodies and clones were used: CD3 (145-2C11), CD4 (RM4-5), CD8 (53-6.7), CD45 (30-F11), FoxP3 (FJK-16s), TCRb (H57-597), CD25 (PC61.5), CTLA4 (UC10-4B9), ICOS (C398.4A), Ki67 (B56), CD44 (IM7), CD69 (H1.213), IL-17 (TC11-18H10.1), IFNγ (XMG1.2), TNFα (MP6-XT22). Samples were run on a Fortessa analyzer (BD Biosciences) for conventional flow cytometry or an Aurora analyzer (Cytek) for spectral flow cytometry in the UCSF Flow Cytometry Core. Data was collected using FACS Diva software (BD Biosciences). Data were analyzed using FlowJo software (FlowJo, LLC). Dead cells and doublet cell populations were excluded, followed by pre-gating on CD45^+^ populations for immune cell analysis. Lymphoid cells were gated as TCRβ^+^CD3^+^ T cells, CD3^+^CD8^+^ T cells (CD8), CD3^+^CD4^+^CD25^−^Foxp3^−^ T effector cells (Teff), and CD3^+^CD4^+^CD25^+^Foxp3^+^ regulatory T cells (Tregs). For human Tregs, expression of layilin was quantified based on isotype control antibody or secondary control.

### RNA-Sequencing Analysis

For human Tregs, cells were sorted based on layilin expression and put in culture for 24h in the presence of anti-CD3/ antiCD28 stimulation. After 24h, cells have been harvested and spun down to obtain a pellet, followed by a flash freezing of the subsequent. Samples have been shipped to Novogene for RNA extraction, library preparation, and sequencing.

For the imiquimod mouse model, Treg and Teff cells were isolated respectively by gating on live CD45^+^CD3^+^CD4^+^CD8^-^CD25^hi^CD62L^hi^GFP^+^ and CD45^+^CD3^+^CD4^+^CD8^-^CD25^lo^CD62L^hi^GFP^-^cells. Cells were sorted into lysis buffer from the kit and frozen on dry ice before being processed using the SMART-Seq® v4 Ultra® Low Input RNA Kit (Takara Bio) according to the manufacture’s protocol. 100pg of amplified cDNA obtained was used to generate a library using Nextera XT DNA preparation kit (Illumina). The quality of cDNA and library products was checked and quantified using an Agilent High Sensitivity DNA chip. An equal quantity of libraries was indexed and pooled together before being sent to Novogene.

We used Novaseq6000 PE150 to a 30M read depth for all sequencing. Reads were aligned to Ensembl GRCm39 (for mouse) or GRCh38 (for human) reference genome using Kallisto. The R/Bioconductor package Limma voom was used to determine differential expression (*47*, *48*). Sequencing data will be uploaded to the National Center for Biotechnology Information Gene Expression Omnibus (GEO) upon acceptance of the manuscript.

### *In vitro* human Treg suppression assay

Before plating, the responder cells (CD4^+^ T effectors and CD8^+^ T cells) were stained with cell trace violet (CTV, ThermoFisher) at 7 million/ml concentration and CTV diluted at 1:8000 in PBS. After a 30-minute incubation in the dark at 37 ° C, cells were washed and resuspended at a concentration of 5×10^5^ cells/ml. 50K of responder cells were plated in 96 well U-bottom plates, and appropriate cell numbers equivalent to each ratio of Tregs CRISPR were added to the plate, as well as anti-CD3/anti-CD28 stimulation. After a three-day incubation, the proliferation of responder cells was measured using CTV quantification and flow cytometry.

### Imiquimod model

Mice were treated for 5 days with tamoxifen to induce Cre recombinase before the start of the treatment. On the last day of tamoxifen treatment, the back skin of the mice were shaved. Two days later, we measured the clinical score of mice to establish a baseline with the following readouts: erythema, scaling, skin lesions, and weight. We then started the imiquimod 5% cream (Taro) application for six consecutive days. On day 7 (for flow cytometry) or day 5 (for bulk RNAseq), mice were sacrificed, and skin and sdLN were harvested. Samples were stained and analyzed by flow cytometry.

### LFA-1 activation assay

100,000 cells were plated in a 96-well U-plate for 24 hours in combination with appropriate treatment. To report LFA-1 activation, cells were stained at 37°C with clone m24 (BioLegend) in an affinity buffer containing 20 mM Hepes, 140 mM NaCl, 1 mM MgCl_2_, 1 mM CaCl_2_, 2 mg/ml glucose, and 0.5% BSA. 2 mM MnCl_2_ was used as a positive control and a combination of 2 mM MnCl_2_ + 2 mM EDTA as negative control. For layilin crosslinking, Layilin antibody was added simultaneously using the following concentrations: 12.5 or 25 ug/ml (Sino Biological).

### *In vitro* human Treg adhesion assay

The day before, the plate was coated with 2.5ug/ml of collagen IV (C5533, Millipore Sigma) and incubated overnight at 4 degrees. Cells were sorted based on layilin expression and put in culture for 24h in the presence of anti-CD3/ antiCD28 stimulation. After incubation, the plate was washed twice by flipping it to remove any unbound cells. To measure the cells attached, we used Cell Titer Blue (Promega) and read the fluorescence using a plate reader.

### F-actin staining

Cells were sorted based on layilin expression and put in culture for 48h in the presence of anti-CD3/ anti-CD28 stimulation. After 48h, cells were harvested, and a cytospin was performed to transfer cells on a slide. After being fixed for 10 minutes at room temperature using 4% PFA, slides were washed and stained with phalloidin Alexa Fluor 546 (Invitrogen) diluted in a mix of PBS containing 1% BSA and 0.1% saponin for an hour at room temperature protected from light. After washing in PBS, slides were mounted with a hard mounting solution containing DAPI (Southern Biotech). Images x100 were obtained using a confocal microscope.

### Quantitative PCR

For assessment of *layilin* gene expression, Tregs and Teffs were sort-purified from skin and sdLNs of FOXP3-ERT2-Cre; Layn^fl/fl^ and FOXP3-ERT2-Cre mice. RNA was isolated using a column-based kit (PureLink RNA Mini Kit, Thermo Fisher), and then transcribed (iScript cDNA Synthesis Kit, Bio-Rad). Expression of *layilin* was determined using a SYBR Green assay (SSo Advanced Universal SYBR Green kit; Biorad). The cycle number of duplicate or triplicate samples was normalized to the expression of the endogenous control Rsp17. Primer sequences used are for Rsp17: For: 5’ - CGCCATTATCCCCAGCAAG - 3’; Rev 5’ - TGTCGGGATCCACCTCAATG - 3’, for Layn: For: 5’ - TCCATGACGCCTTTCAAAGAC - 3’; Rev 5’-AG-GCTGTGTTATTGCTCTGTTTC-3’. Data are presented as relative arbitrary units (AU).

### Statistical Analyses

Statistical analyses were performed with Prism software package version 10 (GraphPad). P values were calculated using two-tailed unpaired or paired Student’s *t*-test for comparison between two groups, one or two-way ANOVA, followed by the appropriate posthoc test for comparison including more than two groups and/or more than one condition. Pilot experiments were used to determine the sample size for animal experiments. No animals were excluded from the analysis unless there were technical errors. Mice were age- and gender-matched and randomly assigned into experimental groups. Appropriate statistical analyses were applied, assuming a normal sample distribution. All *in vivo* mouse experiments were conducted with at least 2-3 independent animal cohorts. RNA-Seq experiments were conducted using 4-5 biological samples (as indicated in figure legends). Data are mean ± S.E.M. P values correlate with symbols as follows: ns = not significant, p>0.05, *p<0.05, **p<0.01, ***p<0.001, ****p<0.0001.

## Acknowledgments

The authors thank the patient donors for providing tissue samples for this study. We thank Clare Abram and Cuyler Luck for assistance with Western blotting. Flow cytometry data were generated in the UCSF Parnassus Flow CoLab (RRID: SCR_018206), supported in part by Grant NIH P30 DK063720 and by the NIH S10 Instrumentation Grant S10 1S10OD026940-01 and S10 1S10OD021822-0. This work was primarily supported by TRex Bio Inc.

## Contribution

Conceptualization: V.G and M.D.R. Methodology: V.G. and M.D.R. Investigation: V.G., C.E.M, J.N.C., S.C., M.K, J.V, H.A.T, M.P., M.M.L, K.R, and A.Z. Resources: H.H., E.K., and I.N. provide human skin samples. S.J.L., K.S, and D.R.M provide confocal microscope and expertise for actin polymerization quantification. Data curation: V.G., S.C., J.V. Writing—original draft: V.G. Writing—editing and revision: V.G., K.R, A.A.Z and M.D.R. Supervision: M.D.R.

## Data and materials availability

Upon acceptance: scRNAseq data are available on GEO with accession numbers XXX. Bulk RNAseq data are available on GEO with accession numbers XXX (for humans) and XXX (for mice). All correspondence and requests for materials and coding scripts can be made to the corresponding author, M.D.R. All other data needed to evaluate the conclusions of this study are present in the paper.

---

**Figure S1.**

**A.** Volcano plot comparing Treg cluster from psoriasis to Treg cluster from healthy skin. **B.** Quadrant representation of results from pseudo bulk RNAseq of Treg cluster subclustered in *layilin* expressing and non-expressing Tregs. Red dots represent differentially expressed genes; black dots are not differentially expressed ones. The left upper quadrant corresponds to genes enriched only in psoriatic *layilin* expressing Tregs, the right bottom quadrant are the ones only enriched in *layilin* expressing Tregs from healthy skin, and upper right quadrant corresponds to genes enriched in *layilin* expressing Tregs and common to both conditions.

**Figure S2.**

**A.** qRT-PCR of *layilin* from sorted Tregs and Teff cells isolated respectively by gating on live CD45^+^CD3^+^CD4^+^CD8^-^CD25^hi^CD62L^hi^GFP^+^ and CD45^+^CD3^+^CD4^+^CD8^-^CD25^lo^CD62L^hi^GFP^-^. Tregs and Teff have been isolated from back skin of FOXP3-ERT2-Cre; Layn^fl/fl^ or FOXP3-ERT2Cre mice after four days of treatment. Each bar corresponds to 5 mice mixed to create a representative sample for each group. **B.** MFI of cells Teff, CD8, and Treg either IFNɣ+, IL17+, or TNFα^+^. n =7 and 6 mice per group from three independent experiments. Significance was defined by a multiple-paired t-test. **C.** MFI of Tregs activation markers gated on Live CD3^+^TCRβ^+^ CD4^+^FOXP3^+^ Tregs. n =7 and 6 mice per group from three independent experiments, except ICOS and CTLA from 2 mice and one experiment. Significance was defined by a multiple-paired t-test. **D.** Quality control of samples repartition used for bulk RNAseq in figure 5. **E.** Quality check of bulk RNAseq representing total of counts for each sample. In F and G, n = 5 mice per group from two independent experiments.

**Figure S3.**

**A.** Quality check of bulk RNAseq representing total of counts for each sample (left panel) and *layn* count for each sample demonstrating CRISPR efficiency (right panel). n = 4 paired donors for each group. **B.** On the left panel, we represent the frequency of total m24+ Tregs gated on Live CD8-CD4+CD25+FOXP3+ cells after three days of culture with or without anti-CD3/anti-CD28 stimulation and in the presence of a layilin crosslinking antibody as described on the x-axis for 20 minutes during LFA-1 activation staining. The right panel represents the frequency of CD18^+^ Tregs. n = 3 donors. Significance was defined by one-way ANOVA followed by Dunnet’s multiple comparisons test. *p < 0.05, ***p<0.001. **C.** Frequency of LAYN^+^ Tregs gated on Live CD8^-^ CD4^+^CD25^+^FOXP3^+^ cells to measure CRISPR efficiency after 24 hours of culture in a non-coated plate or collagen IV-coated plate. n = 6 donors from two independent experiments.

